# Met regulates endoderm migration in zebrafish

**DOI:** 10.64898/2026.05.01.722050

**Authors:** Po-Shu Tu, Aaliyah M. Ruiz-Corral, Stephanie Woo, Stefan C. Materna

## Abstract

Cells can employ different modes of migration, switching between them depending on context. However, how migration modes are determined remains incompletely understood. The mode of migration depends not only on external signals and guidance cues but also on which cell surface receptors a cell expresses. Receptor tyrosine kinases (RTKs) are central mediators of many processes including cell migration, yet whether RTK signaling mediates shifts in migratory behavior in vivo remains unclear. Here, we show that the RTK Met promotes persistent, directional migration of endodermal cells during gastrulation in zebrafish. *met* is broadly expressed across migrating endoderm, and pharmacological inhibition or genetic loss of its function delays endoderm convergence. Quantitative live imaging and cell tracking reveal that loss of Met reduces displacement and persistence without substantially affecting velocity, indicating that Met promotes directional migration rather than motility per se. Although Met is canonically activated by hepatocyte growth factor (Hgf), expression of *hgfa* and *hgfb* during gastrulation is spatially restricted and temporally limited. Consistent with this, genetic loss of Hgf function indicates that it is dispensable for endoderm convergence and migration. Together, these findings identify Met as a regulator of migratory persistence during endoderm convergence and suggest a ligand-independent mode of RTK function in the regulation of cell behavior during development.

**Highlights:**

- Met promotes directional migration of endoderm cells during convergence.
- Loss of Met delays convergence by reducing cell displacement and persistence without affecting velocity.
- Hgf signaling is dispensable for endoderm convergence despite being the canonical Met ligand.

## INTRODUCTION

Cell migration can take many forms — from collective to single-celled, or from directed and persistent to random and exploratory. Different cell types can employ different modes of migration, and the same cell can switch modes depending on context (Petrie et al., 2009). A cell’s choice of migration mode depends not only on external signals and guidance cues but also on the cell’s intrinsic state, including the exact repertoire of cell surface receptors it is expressing at any given moment.

During early zebrafish development, endoderm cells undergo a pronounced transition from low-persistence dispersal to coordinated migration toward the midline, becoming increasingly directional by mid-gastrulation (Pézeron et al., 2008; Woo et al., 2012). This transition represents a key step in endoderm morphogenesis; however, the signaling mechanisms that enable this behavioral shift remain incompletely understood.

The receptor tyrosine kinase Met is activated by hepatocyte growth factor (Hgf) (Bottaro et al., 1991; Naldini et al., 1991) and a well-established regulator of cell motility and migration (Stanislovas and Kermorgant, 2022; Trusolino et al., 2010). During vertebrate development, Met signaling is required for migration of multiple cell populations, including neurons, muscle precursors (Elsen et al., 2009; Isabella et al., 2020; Lin et al., 2006; Talbot et al., 2019; Tsai et al., 2015). In zebrafish, *met* expression is initiated following the onset of gastrulation and becomes broadly expressed across migrating endodermal cells during mid-gastrulation (Latimer and Jessen, 2008). Its function in endoderm migration, however, has not been examined in detail. Moreover, the spatially restricted expression of its canonical ligand Hgf during gastrulation (Latimer and Jessen, 2008) raises questions about how Met signaling is activated in this context.

Migration of zebrafish endoderm is regulated by multiple signaling pathways. The TGF-β ligand Nodal not only specifies endoderm, but also promotes dynamic migration of endodermal cells, in part through activation of cytoskeletal regulators (Woo et al., 2012). In addition, signaling by the chemokine receptor Cxcr4 and its ligand Cxcl12b coordinates endoderm and mesoderm movements and contributes to directional migration toward the midline (Chang et al., 2025; Mizoguchi et al., 2008; Nair and Schilling, 2008). However, disruption of these pathways delays convergence but does not completely abolish motility, indicating that distinct components of migratory behavior—such as speed, displacement, and persistence—are differentially regulated and that additional mechanisms contribute to the establishment of persistent, directional migration.

Here, we report that *met* is ubiquitously expressed in migrating endodermal cells during gastrulation and promotes efficient endoderm convergence. Loss of Met reduces cell displacement and persistence without substantially affecting velocity, indicating that Met enables persistent directional migration rather than motility per se. Although Met is canonically activated by Hgf, the expression patterns of *hgfa* and *hgfb* during gastrulation are spatially restricted and not consistent with a role in endoderm migration. Accordingly, genetic evidence indicates that Hgf signaling is dispensable for this process. Together, these findings identify Met as a regulator of migratory persistence during endoderm convergence and reveal a ligand-independent role for Met signaling in this context.

## RESULTS

### *met* is uniformly expressed throughout the migrating endoderm

While previous studies have reported *met* expression in presumptive endoderm (Latimer and Jessen, 2008), the extent to which *met* is uniformly expressed across migrating endodermal cells has not been resolved. Notably, *met* is not expressed during early gastrulation stages (Latimer and Jessen, 2008), suggesting that its expression is temporally regulated. To address this, we visualized its spatial distribution using hybridization chain reaction (HCR) RNA fluorescent in situ hybridization (RNA-FISH) in *Tg(sox17:GFP)^s870^*embryos, which label endodermal cells (Mizoguchi et al., 2008). At 10 hours post-fertilization (hpf), i.e., the end of gastrulation, *met* transcripts were detected throughout the endoderm (Fig. 1A–A′′). Notably, we observed complete concordance between *met* signal and *sox17:*GFP within the endoderm, indicating that *met* is expressed across the entire population of migrating endodermal cells (Fig. 1A′′). Although *Tg(sox17:GFP)^s870^* also labels dorsal forerunner cells (DFCs), these cells lacked appreciable *met* expression, consistent with previous reports (asterisks, Fig. 1A–B) (Latimer and Jessen, 2008). Thus, within the *sox17*-lineage, *met* expression is specific to endoderm. In addition, we observed *met* expression in a restricted anterior domain outside the endoderm (arrowhead, Fig. 1A), consistent with anterior neuroectoderm as previously described (Elsen et al., 2009).

**Figure 1.**
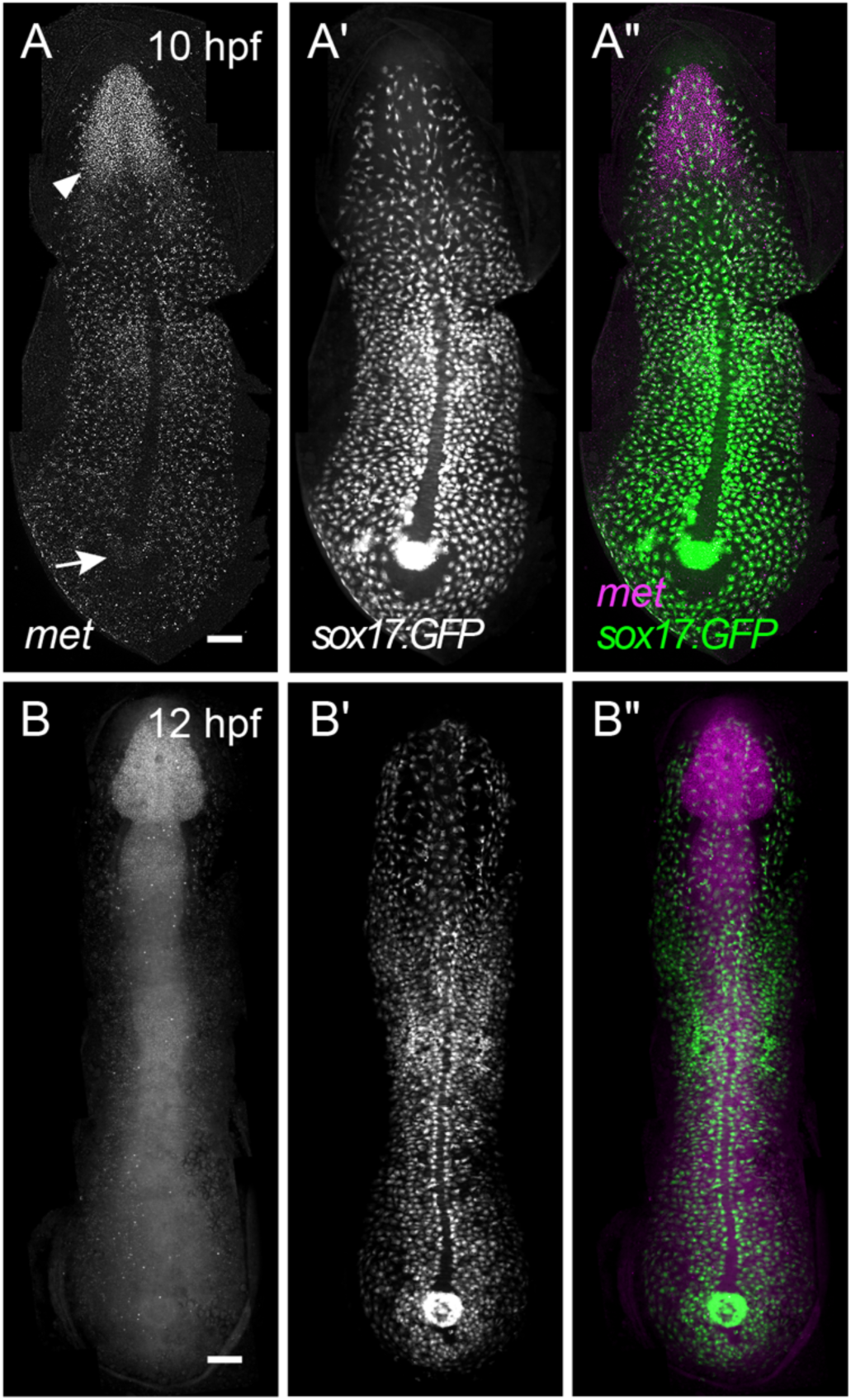
*met* is uniformly expressed throughout endodermal cells. Representative images of HCR RNA-FISH for *met* in flat-mounted *Tg(sox17:GFP)* embryos at 10 hpf (A–A′′) and 12 hpf (B–B′′). Panels show *met* signal (A, B and magenta in A′′, B′′) and GFP (A′, B′ and green in A′′, B′′). Arrow, Kupffer’s Vesicle. Arrowhead, anterior neurectoderm (A) and developing hindbrain and anterior neural tube (B). Dorsal views, anterior towards the top. Scale bars, 100 μm.

This primarily endodermal expression pattern persisted until 12 hpf (Fig. 1B–B′′). At this stage, *met* expression in non-endodermal tissues expanded posteriorly (Fig. 1B), consistent with its reported expression in forebrain, dorsal midbrain, and hindbrain, forming a gradient that tapers in the posterior neural tube (Elsen et al., 2009). Collectively, these data indicate that while *met* is mostly restricted to the endoderm during gastrulation, its expression in non-endodermal tissues expands following gastrulation.

### Inhibition of Met delays endoderm convergence

From mid- to late-gastrulation, endodermal cells transition from dispersal to more directional migration and eventually converge on the midline (Pézeron et al., 2008; Williams and Solnica-Krezel, 2020; Woo et al., 2012). Notably, this shift in migratory behavior coincides with peak *met* expression in the endoderm, suggesting that Met signaling may promote efficient directional migration during convergence.

Activation of Met depends on ligand-induced dimerization and autophosphorylation, which enables recruitment of downstream signaling components. To test whether Met activity is required for endoderm convergence, we used two established Met inhibitors that block Met phosphorylation and downstream signaling, INCB28060 (Capmatinib) (Liu et al., 2011) and SGX-523 (Buchanan et al., 2009). *Tg(sox17:GFP)^s870^*embryos were treated with either Met inhibitor or vehicle control starting at 3.5 hpf. To confirm inhibitor efficacy, we assessed later developmental phenotypes and found that by 3 dpf treated larvae exhibited reduced body length (2.994±0.376 mm vs. 3.429±0.103 mm; *p* < 0.0001, Student’s t-test; Fig. 2 A–B), consistent with the *met^umu7^*mutant line (Nord et al., 2019); treated larvae also had reduced head size, in agreement with prior morpholino-based perturbations of *met* (Elsen et al., 2009), supporting effective Met inhibition. Embryos were then fixed at 10 hpf and imaged. In lateral views (Fig. 2C–E), control embryos exhibited typical ventral-to-dorsal migration, with most endodermal cells positioned dorsally (Fig. 2C). In contrast, Met inhibitor-treated embryos displayed a broader distribution of endodermal cells, with many cells remaining in more ventral positions (Fig. 2D–E), suggesting that convergence is impaired.

**Figure 2.**
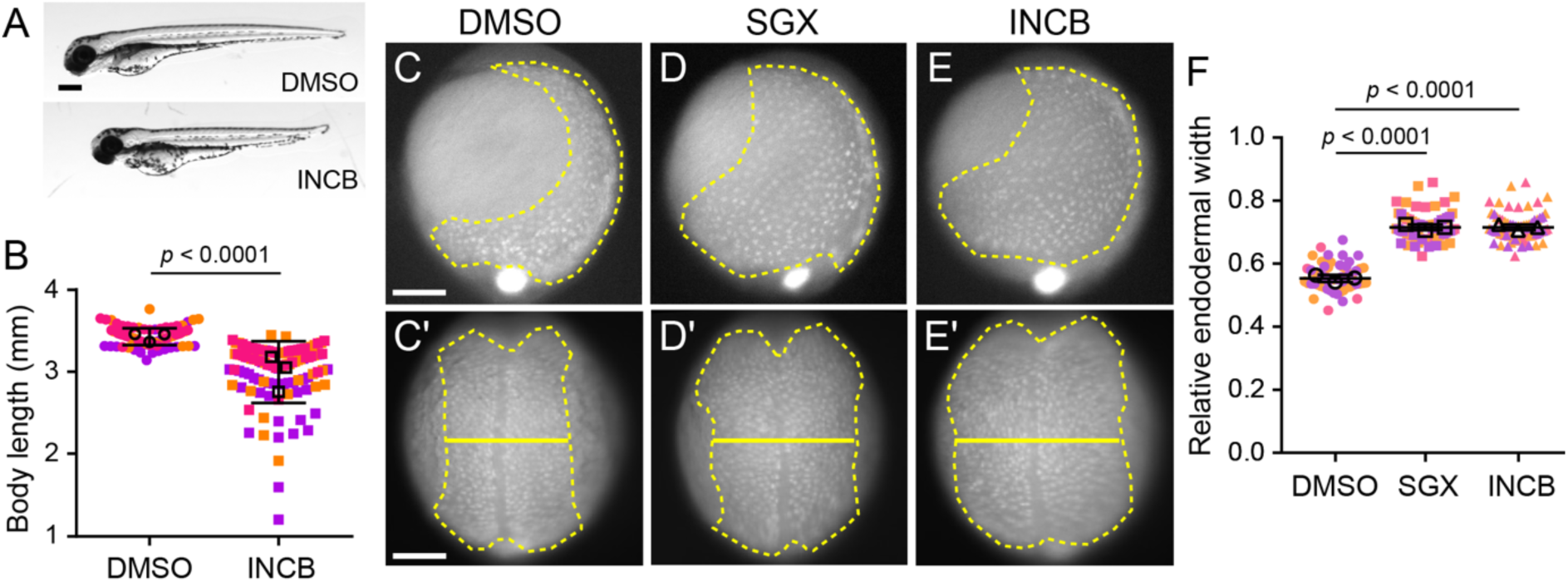
Inhibition of Met delays endoderm convergence. (A) Representative images of 3 dpf larvae treated with DMSO (top) or 30 µM INCB28060 (bottom). Lateral views, anterior to the left. Scale bar, 250 μm. (B) Quantification of body length at 3 dpf of larvae treated with DMSO or 30 µM INCB28060 (INCB). Points represent individual larvae colored coded by batch. Open circles indicate batch means. Error bars indicate standard deviation. *p* values determined by Student’s t-test. DMSO (n = 109) larvae, INCB (n = 109 larvae), from three independent batches each. (C–E′) Representative images of *Tg(sox17:GFP)* embryos treated with DMSO (C, C′), SGX-523 (D, D′), or INCB28060 (E, E′) from 3.5 hpf and imaged at 10 hpf (C–E, lateral view, dorsal to the right) and 11 hpf (C′–E′, dorsal view, anterior towards to the top). Endoderm is outlined (dashed line), and endoderm width is indicated (solid line). Scale bars, 150 μm. (F) Quantification of the relative endodermal width (endoderm/embryo width) at 11 hpf from embryos treated with DMSO, SGX-523 (SGX), and INCB28060 (INCB). Points represent individual larvae colored coded by batch. Open circles indicate batch means. Error bars indicate standard deviation. *p* values determined by Student’s t-test. DMSO (58 embryos), SGX (n = 74 embryos), and INCB (n = 74 embryos) from three independent batches each.

To quantify this phenotype, we measured endodermal width from dorsal views of control and Met inhibitor-treated 10 hpf embryos (Fig. 2C′–E′). To account for variability in embryo size, we calculated the ratio of medial endoderm width to total embryo width (hereafter referred to as relative endodermal width; Fig. 2F). Control embryos exhibited a mean relative endodermal width of 0.557 ± 0.04 (SD), whereas embryos treated with INCB28060 or SGX-523 showed significantly increased values of 0.716 ± 0.04 and 0.715 ± 0.05, respectively (∼28% increase; both *p* < 0.0001, Student’s t-test). Importantly, Met inhibitor-treated embryos did not exhibit delays in epiboly, indicating that the convergence defect is not due to a global developmental delay but is specific to endodermal cell behavior during gastrulation. These results suggest that Met activity is required for efficient endoderm convergence.

### Met inhibition reduces persistence of endodermal cell migration

To further define how Met regulates endoderm convergence, we performed time-lapse confocal imaging of *Tg(sox17:GFP)^s870^* embryos to quantify the dynamics of endodermal cell migration. Given the delay in endoderm convergence observed upon Met inhibition, we sought to determine how Met activity influences migratory behavior at the single-cell level. As above, embryos were treated with Met inhibitor or vehicle control prior to gastrulation (3.5 hpf) and imaged starting at 8.5 hpf over a ∼1.5 hour window that encompasses dorsal convergence of endoderm (Pézeron et al., 2008) (Video 1).

Consistent with our fixed imaging analysis, Met inhibitor-treated embryos exhibited delayed endoderm convergence, with endodermal cells occupying a broader domain at comparable stages (Fig. 3A–B′). Cell-tracking analysis revealed clear differences in migratory behavior (Fig. 3A′′–B′′), with inhibitor-treated cells exhibiting shorter and less directed trajectories. Quantification showed a significant reduction in migratory displacement, velocity, and persistence in Met inhibitor-treated embryos (Fig. 3C–E). Notably, the most pronounced effect was a reduction in migratory persistence defined as the ratio between net and total distance traveled (from 0.718 ± 0.188 to 0.636 ± 0.210, *p* = 0.0189 by Student’s t-test). In contrast, velocity was only slightly affected (from 1.124 ± 0.346 to 1.057 ± 0.321 µm/min, *p* = 0.0116 by Student’s t-test), resulting in an overall decrease in displacement (from 65.26 ± 33.34 to 54.02 ± 30.64 µm, *p* = 0.0025 by Student’s t-test).

**Figure 3.**
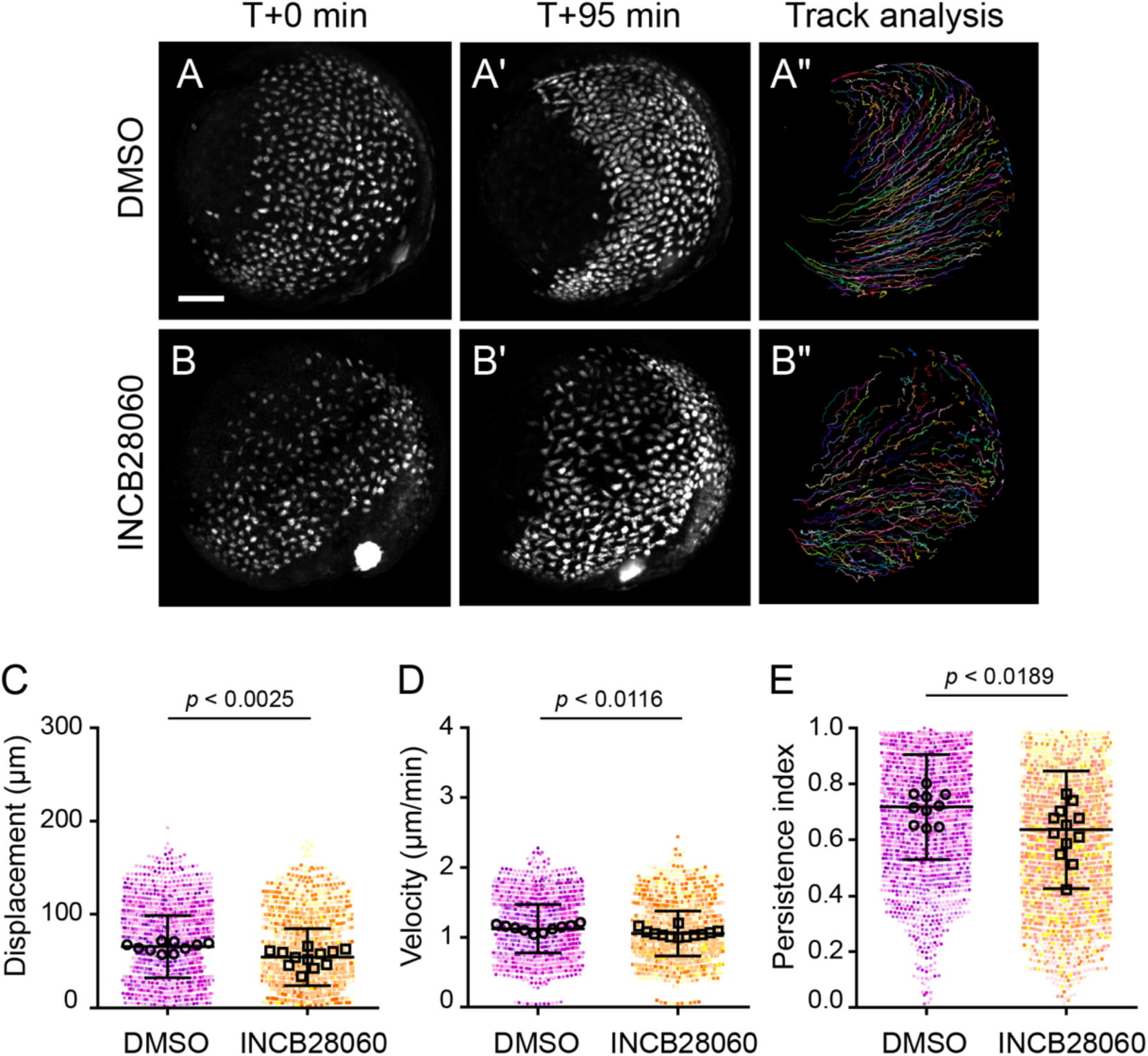
Inhibition of Met reduces displacement and persistence of endodermal cells. (A–B′′) Time-lapse confocal imaging of *Tg(sox17:GFP)* embryos starting at 8.5 hpf. Embryos treated with DMSO (A–A′′) or INCB28060 (B–B′′) are shown at the start (A, B) and end (A′, B′) of each time-lapse along with migration tracks generated by cells throughout the time-lapse (A′′, B′′). Lateral views, dorsal to the right. Scale bar, 100 μm. (C–E) Quantification of migration displacement (C), velocity (D), and persistence (E). Points represent individual cell tracks, color colored by embryo. Open circles depict embryo means. Error bars indicate standard deviation. *p* values determined by one-way ANOVA. DMSO (n = 2742 cells from 10 embryos). INCB28060 (n = 3375 cells from 12 embryos).

Collectively, these results indicate that *met* promotes efficient endoderm convergence primarily by enhancing migratory persistence, which may enable endodermal cells to sustain directional movement toward the midline.

### CRISPR/Cas9-mediated generation of a *met* transcriptional null allele

While pharmacological inhibition strongly suggested a role for Met in endoderm convergence, inhibitor treatments are vulnerable to nonspecific effects. Moreover, previously described *met* alleles, including *met^fh533^* (Isabella et al., 2020), encode truncated proteins that remove key functional domains (e.g., the kinase domain) but retain N-terminal regions, leaving open the possibility of residual activity or compensation via transcriptional adaptation (El-Brolosy et al., 2019). To ensure complete absence of a *met* gene product, we generated *met* loss-of-function mutants using CRISPR/Cas9 (Fig. 4A).

**Figure 4.**
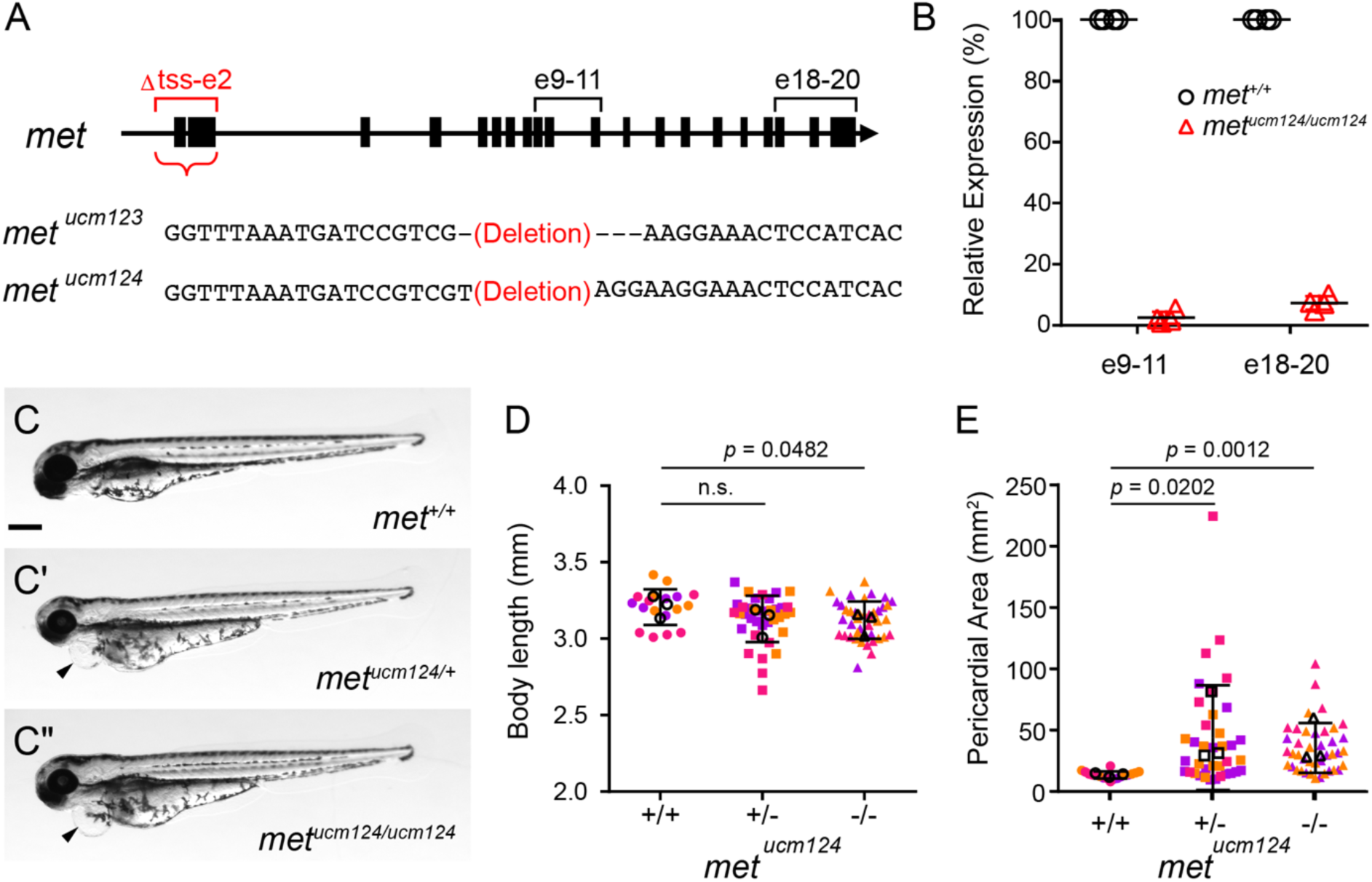
Generation and characterization of met knockout mutants. (A) Schematic of *met* locus and deletion alleles (*met^ucm123^* and *met^ucm124^*), which remove the region spanning the transcription start site through exon 2 (Δtss–e2; red brackets). Black brackets indicate qPCR primers flanking exons 9–11 (e9–11) and exons 18–20 (e18–20). (B) qPCR detection of *met* transcripts from wild-type (*met^+/+^*) or homozygous mutant (*met^ucm124/ucm124^*) embryos at 10 hpf using primers e9–11 and e18–20. Points represent individual embryos. Horizontal bars indicate means. n = 4 embryos per genotype. (C–C′′) Representative images of wild-type (*met^+/+^*, C), heterozygous (*met ^ucm124/+^*, C′) or homozygous mutant (*met^ucm124/ucm124^*, C′′) embryos at 3 dpf. Arrowheads indicate cardiac edema. Scale bar, 250 μm. (D, E) Quantification of body length (D) and pericardial area (E) of wild-type (+/+), heterozygous (+/-), and homozygous mutant (-/-) larvae at 3 dpf. Points represent individual larvae color coded by batch. Open circles indicate batch means. Error bars indicate standard deviation. *P* values determined by one-way ANOVA. n.s., not significant. +/+ (n = 18), +/− (n = 37), and −/− (n = 41), from three independent batches each.

We designed two gRNAs targeting the *met* locus: one upstream of the transcription start site (tss) and one downstream of exon 2, enabling deletion of the promoter and early coding region. This strategy is expected to abolish *met* transcription and, thus, protein production. Screening of F1 animals identified deletions of ∼6 kb (Δtss-e2), resulting in two independent alleles, *met^ucm123^*and *met^ucm124^*. At 10 hpf, when *met* expression is most prominent in the endoderm, we confirmed a near-complete loss of *met* transcripts in *met^ucm124^*embryos by qPCR, indicating effective depletion. Although both alleles survived beyond larval stages, maternal–zygotic mutants could not be recovered due to poor survival and infertility of homozygous adults. These findings are consistent with the previously described *met^fh533^* allele (Isabella et al., 2020), which disrupts the kinase domains of Met. Accordingly, homozygous embryos used in subsequent analyses were generated by intercrossing heterozygous parents. *met^ucm123^* and *met^ucm124^* mutants exhibited indistinguishable phenotypes (results for *met^ucm123^* are summarized in Supp. Fig. 1). For simplicity, we chose to focus on *met^ucm124^* for subsequent characterization.

*met^ucm124^* mutants exhibited no overt morphological defects during embryogenesis. However, by 3 dpf, mutants displayed reduced larval body length (Fig. 4C–D). Wild-type siblings exhibited a mean body length of 3.206 ± 0.117 mm, whereas heterozygous and homozygous mutants measured 3.13 ± 0.152 mm and 3.121 ± 0.121 mm, respectively. This reduction was significant in homozygous mutants compared to wild-type siblings (*p* = 0.0482, Student’s t-test), consistent with both Met inhibitor-treated larvae and the previously described *met^umu7^*mutant (Nord et al., 2019). Similarly, *met^ucm124^* larvae exhibited reduced head size, consistent with the cerebellar defects reported in *met* knockdown embryos (Elsen et al., 2009). Together, these phenotypes are consistent with previously reported *met* perturbations, supporting *met^ucm124^* as a functional loss-of-function allele.

Interestingly, *met^ucm124^* larvae developed pericardial edema by 3 dpf (Fig. 4C–C′′,E). Quantification revealed a significant increase in pericardial area in both heterozygous and homozygous mutants compared to wild-type siblings (*p* = 0.0202 and 0.0012, respectively, one-way ANOVA) (Fig. 4E). The mean pericardial area of wild-type siblings was 13.68 ± 2.94 mm², whereas heterozygous and homozygous mutants exhibited values of 44.05 ± 42.69 mm² and 35.60 ± 20.32 mm², respectively. These findings indicate that loss of *met* function leads to an increase in pericardial area, consistent with pericardial edema.

### Loss of *met* delays endoderm convergence

To determine whether loss of *met* disrupts endoderm convergence, *met^ucm124^* mutant embryos were imaged at 11.5 hpf (Fig. 5A–A′′). Consistent with our observations in Met inhibitor-treated embryos, homozygous *met^ucm124^* mutants exhibited a marked delay in endoderm convergence, with endodermal cells occupying a broader domain (Fig. 5A′′). Notably, heterozygous embryos also displayed an increased relative endodermal width (Fig. 5A′), indicating that *met* is haploinsufficient for proper endoderm convergence. Quantification (Fig. 5C) revealed a significant increase in mean relative endodermal width in both heterozygous (0.525 ± 0.088) and homozygous (0.643 ± 0.097) *met^ucm124^* embryos compared to wild-type siblings (0.459 ± 0.090; *p* = 0.005 and *p* < 0.0001, respectively, one-way ANOVA). Consistent with these findings, *met^fh533^* mutants also exhibited increased relative endodermal width (Fig. 5B–B′′). Heterozygous and homozygous *met^fh533^* embryos displayed mean values of 0.613 ± 0.065 and 0.646 ± 0.068, respectively, compared to 0.521 ± 0.055 in wild-type siblings (both *p* < 0.0001, one-way ANOVA; Fig. 5D). Together, these results demonstrate that loss of *met* function delays endoderm convergence and that this phenotype is consistent across independent alleles affecting either Met expression or kinase activity.

**Fig. 5.**
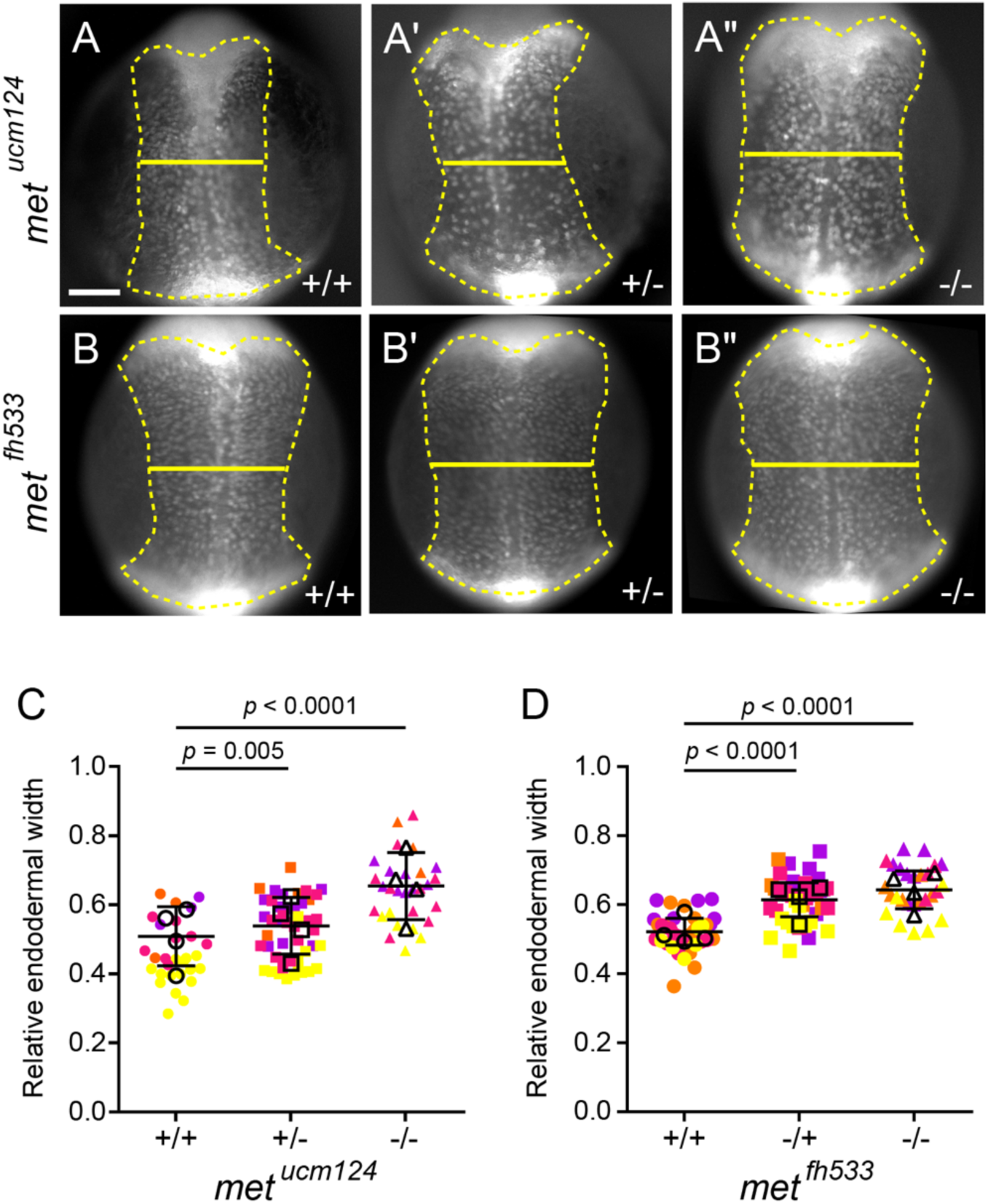
Loss of Met delays endoderm convergence. (A–B′′) Representative images of endoderm labeled with *Tg(sox17:GFP)* in wild-type (A, B), heterozygous (A′ and B′), and homozygous mutant (A′′, B′′) *met^ucm124^*(A–A′′) and *met^fh533^*(B–B′′) embryos imaged at 11.5 hpf (dorsal view, anterior towards the top). Endoderm is outlined (dashed line), and endoderm width is indicated (solid line). Scale bar, 150 μm. (C, D) Quantification of relative endoderm width (endoderm/embryo width) at 11.5 hpf from wild-type, heterozygous, and homozygous mutant *met^ucm124^* (C) and *met^fh533^* (D) embryos. Points represent individual embryos color coded by batch. Open circles indicate batch means. Error bars indicate standard deviation. *p* values determined by one-way ANOVA. *met^ucm124^*: +/+ (n = 29), +/− (n = 48), −/− (n = 29); *met^fh533^*: +/+ (n = 36), +/− (n = 40), −/− (n = 27), from four independent batches each.

### Met promotes persistent, directional migration of endodermal cells

To determine how loss of met delays endoderm convergence, we examined the migratory behavior of endodermal cells by time-lapse imaging. As above, imaging was performed from 8.5 hpf and spanned mid- to late-gastrulation (Fig. 6A–C′′, Video 2). Consistent with our inhibitor experiments, *met^ucm124^* mutants exhibited delayed endoderm convergence (Fig. 6C). Cell-tracking analysis revealed qualitatively shorter and more tortuous migratory trajectories in both heterozygous and homozygous mutants compared to controls (Fig. 6A′′–C′′).

**Figure 6.**
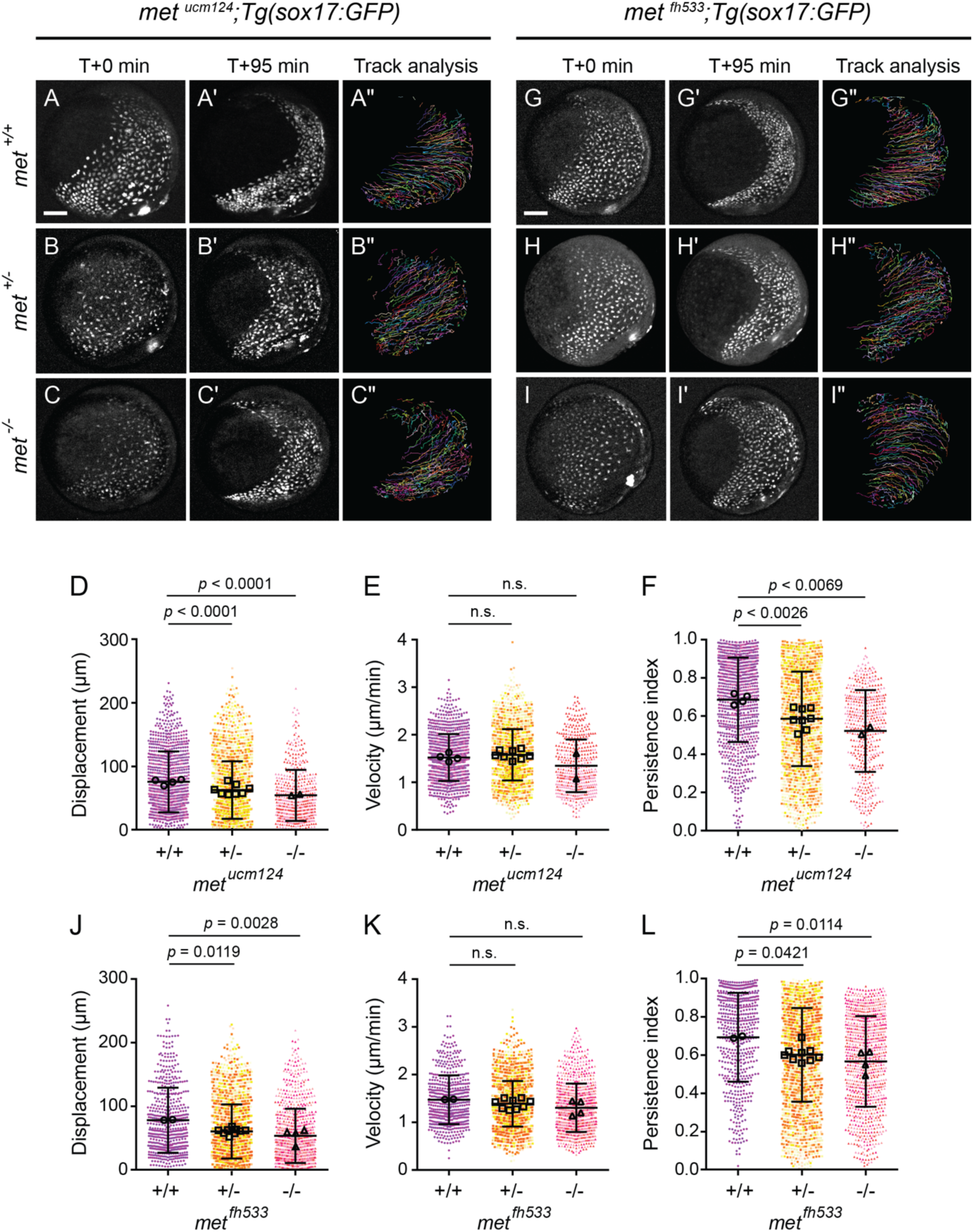
Loss of Met reduces displacement and persistence of endodermal cells. (A–C′′) Time-lapse confocal imaging of endoderm labeled with *Tg(sox17:GFP)* in wild-type (A–A′′), heterozygous (B–B′′), and homozygous mutant (C–C′′) *met^ucm124^* embryos starting at 8.5 hpf. Embryos are shown at the start (A, B, C) and end (A′, B′, C′) of each time-lapse along with migration tracks generated by cells throughout the time-lapse (A′′, B′′, C′′). Lateral views, dorsal to the right. Scale bar, 100 μm. (D–F) Quantification of migration displacement (D), velocity (E), and persistence (F) from wild-type, heterozygous, and homozygous mutant *met^ucm124^*embryos. Points represent individual cell tracks, color coded by embryo. Open circles depict embryo means. Error bars indicate standard deviation. *p* values determined by one-way ANOVA. (G–I′′) Time-lapse confocal imaging of endoderm labeled with *Tg(sox17:GFP)* in wild-type (G–G′′), heterozygous (H–H′′), and homozygous mutant (I–I′′) *met^fh533^* embryo starting at 8.5. Embryos are shown at the start (G, H, I) and end (G′, H′, I′) of each time-lapse along with migration tracks generated by cells throughout the time-lapse (G′′, H′′, I′′). Lateral views, dorsal to the right. Scale bar, 100 μm. (J–L) Quantification of migration displacement (J), velocity (K), and persistence (L) from wild-type, heterozygous, and homozygous mutant *met^fh533^* embryos. Points represent individual cell tracks, color colored by embryo. Open circles depict embryo means. Error bars indicate standard deviation. *p* values determined by one-way ANOVA. *met^ucm124^*: +/+ (966 cells from 4 embryos), +/− (1995 cells from 8 embryos), −/− (477 cells from 2 embryos); *met^fh533^*: +/+ (577 cells from 2 embryos), +/− (2366 cells from 9 embryos), −/− (906 cells from 4 embryos).

To quantify these changes in cell migration, we measured migration displacement, persistence, and instantaneous velocity in wild-type, heterozygous, and homozygous *met^ucm124^* embryos and compared by one-way ANOVA (Fig. 6D–F). Mean displacement was significantly reduced from 76.06 ± 47.84 µm in wild-type siblings to 63.12 ± 45.34 µm (*p* < 0.001) in heterozygotes and 54.86 ± 40.13 µm (*p* < 0.001) in homozygous mutants. Similarly, persistence significantly decreased from 0.686 ± 0.221 in wild-type embryos to 0.586 ± 0.248 (*p* = 0.0026) in heterozygotes and 0.522 ± 0.214 (*p* = 0.0069) in homozygous mutants. In contrast, instantaneous velocity remained comparable across wild-type (1.521 ± 0.493 µm/min), heterozygous (1.579 ± 0.541 µm/min, *p* = 0.75), and homozygous mutant (1.349 ± 0.554 µm/min, *p* = 0.285) embryos.

Consistent with these findings, *met^fh533^* mutants also exhibited reduced displacement and persistence without a corresponding change in velocity (Fig. 6G–I′′, Video 3). Mean displacement significantly decreased from 78.32 ± 51.14 µm in wild-type embryos to 60.62 ± 42.41 µm (*p* = 0. 0119) and 53.79 ± 42.57µm (*p* = 0. 0028) in heterozygous and homozygous mutants, respectively, while persistence decreased from 0.693 ± 0.232 to 0.602 ± 0.245 (*p* = 0. 0421) and 0.567 ± 0.237 (*p* = 0. 0114). Instantaneous velocity remained comparable across genotypes (1.474±0.511 µm/min in wild-type embryos vs. 1.391±0.479 µm/min in heterozygous embryos, *p* = 0.5063, and 1.304±0.506 µm/min in homozygous embryos, *p* = 0.1822).

Together, these results indicate that *met* regulates endodermal cell migration by promoting migratory persistence and, consequently, efficient displacement, without significantly affecting velocity. Thus, Met primarily supports directional migration rather than overall motility during endoderm convergence.

### *hgf* expression is limited during endoderm convergence

It is well established that Met activation depends on Hgf binding, which induces receptor dimerization and autophosphorylation to initiate downstream signaling (Niemann, 2013; Uchikawa et al., 2021). Given the role of Met in regulating endoderm migration and convergence, we next examined whether Hgf is also required for this process.

Because of a teleost-specific genome duplication, the zebrafish genome contains two *hgf* genes, *hgfa* and *hgfb*, whose expression during embryogenesis has been characterized. *hgfa* expression is first detected toward the end of gastrulation and is restricted to deep anterior cells and adaxial cells (Latimer and Jessen, 2008); at later stages, it is observed in somites, fin buds, and the swim bladder (Anderson et al., 2013; Elsen et al., 2009; Latimer and Jessen, 2008). In contrast, *hgfb* expression initiates later, during somitogenesis, and is localized to the otic vesicle, brain ventricles, and periocular region (Anderson et al., 2013; Elsen et al., 2009; Latimer and Jessen, 2008). Functional studies have shown that perturbation of either ligand impairs processes such as motor neuron and myogenic precursor migration, with more severe phenotypes observed upon combined loss of *hgfa* and *hgfb* (Anderson et al., 2013; Elsen et al., 2009; Isabella et al., 2020; Talbot et al., 2019). Notably, these reported expression patterns do not indicate broad *hgfa* or *hgfb* expression during gastrulation or in proximity to migrating endodermal cells.

To better define the temporal expression of *hgfa* and *hgfb*, we performed qPCR across early developmental stages. *hgfa* transcripts are maternally deposited but decline to near-undetectable levels by early gastrulation, preceding their zygotic upregulation toward the end of gastrulation (Fig. 7A), whereas *hgfb* expression remains minimal throughout this period. Both *hgfa* and *hgfb* increase during somitogenesis, consistent with prior studies (Elsen et al., 2009; Latimer and Jessen, 2008). To assess the spatial distribution of *hgfa* transcripts during gastrulation, we performed HCR RNA-FISH on 10 hpf embryos (Fig. 7B–B′′). Consistent with previous reports (Latimer and Jessen, 2008), *hgfa* expression was restricted to the anterior region of the embryo and to a small adaxial domain along the trunk (arrowheads, Fig. 7B), rather than broadly distributed. Together, these data indicate that, in contrast to the broad expression of *met* in migrating endodermal cells, Hgf ligand availability during gastrulation is limited, with *hgfa* present at low levels in spatially restricted domains and *hgfb* largely absent. These results suggest that neither *hgfa* nor *hgfb* appears to be expressed at the appropriate time or in a sufficiently broad domain to activate Met during endoderm convergence.

**Figure 7.**
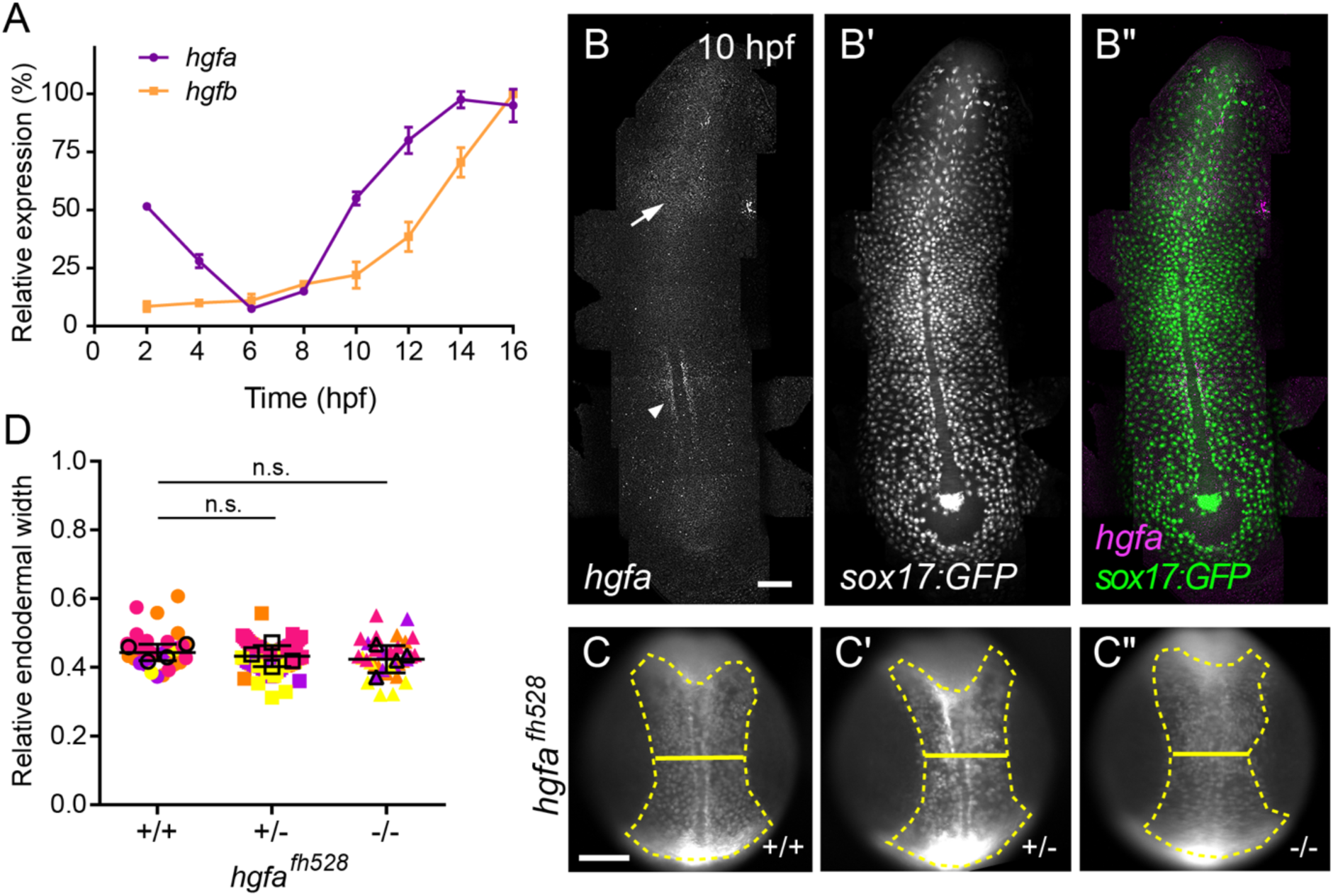
*hgfa* is dispensable for endoderm convergence. (A) qPCR profiling of *hgfa* and *hgfb* expression during early development (from 2 to16 hpf). (B–B′′) Representative images of HCR RNA-FISH for *hgfa* in *Tg(sox17:GFP)* embryos at 10 hpf. Panels show *hgf* signal (B and magenta in B′′), and *sox17:GFP* (B′ and green in B′′). Arrowhead, paraxial mesoderm. Arrow, deep anterior cells. Dorsal views, anterior towards the top. (C–C′′) Endoderm labeled with *Tg(sox17:GFP)* from wild-type (C), heterozygous (C′), and homozygous mutant (C′′) *hgfa^fh528^* embryos at 11.5 hpf. Endoderm is outlined (dashed line), and endoderm width is indicated (solid line). Dorsal views, anterior towards the top. Scale bar, 150 μm. (D) Quantification of relative endoderm width (endoderm/embryo width) at 11.5 hpf from wild-type, heterozygous, and homozygous mutant *hgfa^fh528^*embryos. Points represent individual embryos color coded by batch. Open circles indicate batch means. Error bars indicate standard deviation. *p* values determined by one-way ANOVA. n.s., not significant. +/+ (n = 32), +/− (n = 69), −/− (n = 33), from four independent batches each.

### *hgfa* is dispensable for endoderm convergence

Although *hgfa* expression during gastrulation is low and spatially restricted, it could nonetheless activate Met signaling. If Hgfa-mediated activation of Met is required, loss of *hgfa* function would be expected to delay endoderm convergence.

To test this, we analyzed the previously described *hgfa^fh528^*mutant (Isabella et al., 2020), which carries a 79-bp deletion in exon 6 of the *hgfa* locus that introduces a premature stop codon and results in loss of function. As above, embryos were raised and endoderm convergence was assessed at 11.5 hpf by measuring relative endodermal width (Fig. 7C–C′′). In contrast to Met loss-of-function embryos, *hgfa^fh528^* mutants exhibited normal endoderm convergence. Mean relative endodermal width was comparable across wild-type (0.452 ± 0.0526), heterozygous (0.432 ± 0.0426, *p* = 0.0949), and homozygous (0.427 ± 0.0543, *p* = 0.0669) siblings (Fig. 7C–C′′, 7D).

To determine whether loss of *hgfa* affects endodermal cell migration, we examined the migratory behavior of endodermal cells by time-lapse imaging of *hgfa^fh528^* mutant siblings. As above, imaging started at 8.5 hpf and spanned mid- to late-gastrulation (Fig. 8A–C′′, Video 4). In contrast to *met* mutants, loss of *hgfa* did not alter the overall trajectories of endodermal cells (Fig. 8A′′–C′′). Quantitative analysis revealed no significant differences in migration behaviors (one-way ANOVA; Fig. 8D–F). Mean displacement in wild-type embryos was 67.81 ± 42.14 µm, compared to 66.51 ± 47.82 µm in heterozygotes (*p* = 0.9907), and 65.69 ± 44.24 µm in homozygous mutants (*p* = 0.8033). Mean instantaneous velocity was 1.281 ± 0.436 µm/min in wild-type embryos, compared to 1.337 ± 0.480 µm/min in heterozygotes (*p* = 0.5668) and 1.363 ± 0.484 µm/min in homozygous mutants (*p* = 0.2354). Finally, mean persistence was 0.702 ± 0.235 in wild-type embryos, compared to 0.665 ± 0.250 in heterozygotes (*p* = 0.612) and 0.659 ± 0.243 in homozygous mutants (*p* = 0.3161). Together, these results indicate that, unlike *met*, loss of *hgfa* does not affect endoderm convergence or migration.

**Fig. 8.**
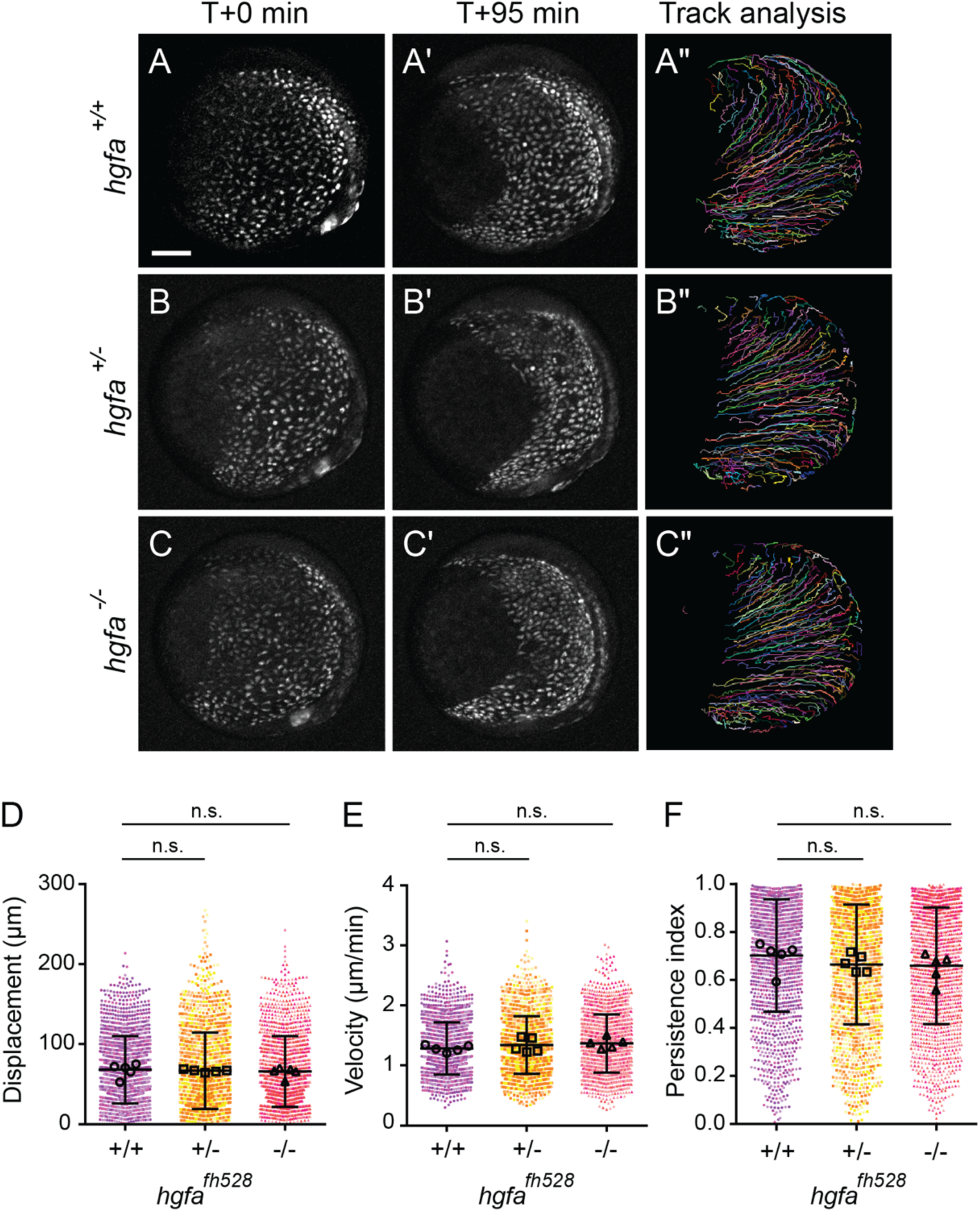
Loss of *hgfa* does not affect displacement or persistence of endodermal cells. (A–C′′) Time-lapse confocal imaging of endoderm labeled with *Tg(sox17:GFP)* in wild-type (A–A′′), heterozygous (B–B′′), and homozygous mutant (C–C′′) *hgfa^fh528^* embryos starting at 8.5. Embryos are shown at the start (A, B, C) and end (A′, B′, C′) of each time-lapse along with migration tracks generated by cells throughout the time-lapse (A′′, B′′, C′′). Lateral views, dorsal to the right. Scale bar, 100 μm. (D–F) Quantification of migration displacement (C), velocity (D), and persistence (E) from wild-type, heterozygous, and homozygous mutant *hgfa^fh528^*embryos. Points represent individual cell tracks, color colored by embryo. Open circles depict embryo means. Error bars indicate standard deviation. *p* values determined by one-way ANOVA. *hgfa^fh528^*: +/+ (1533 cells from 5 embryos), +/− (1571 cells from 5 embryos), − /− (1562 cells from 5 embryos).

### *hgfb* does not compensate for loss of *hgfa*

Genetic compensation can mask mutant phenotypes through transcriptional adaptation (El-Brolosy et al., 2019). We therefore asked whether *hgfb* might be upregulated in *hgfa^fh528^* mutants, potentially compensating for the loss of *hgfa*. qPCR analysis at 10 hpf revealed that *hgfb* levels were unchanged across genotypes (Fig. 9A). These data indicate that *hgfb* expression levels do not increase in response to loss of *hgfa*.

**Figure 9.**
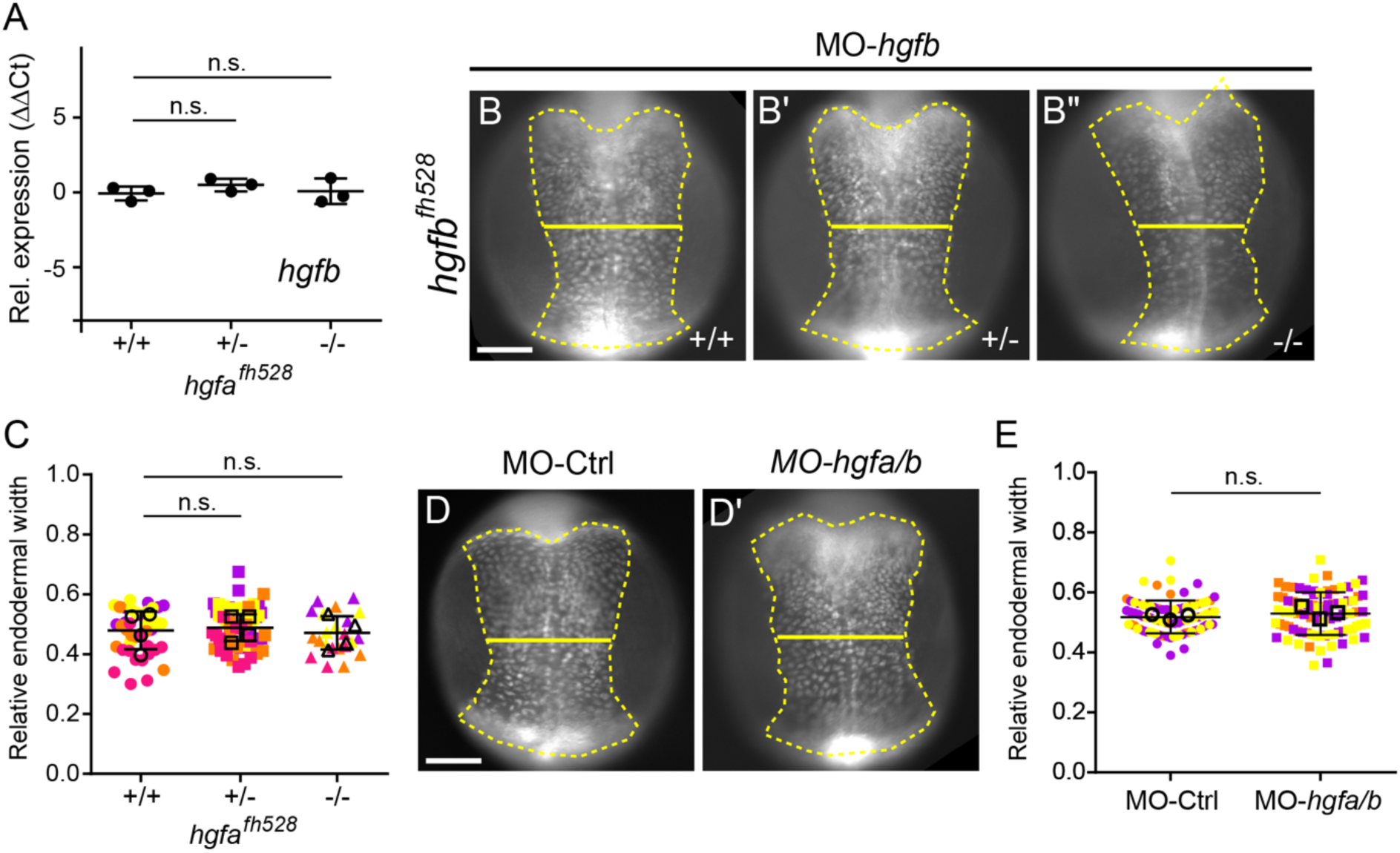
*hgfb* does not compensate for loss of *hgfa* during endoderm convergence. (A) qPCR quantification of expression of *hgfb* in *hgfa^fh528^* mutants relative to wild type siblings. Points represent individual embryos. Horizontal bars indicate means. Error bars indicate standard deviation. *p* values determined by one-way ANOVA. n = 3 embryos per genotype. n.s., not significant. (B–B′′) Endoderm labeled with *Tg(sox17:GFP)* from wild-type (C), heterozygous (C′), and homozygous mutant (C′′) *hgfa^fh528^* embryos injected with *hgfb* morpholino (MO-*hgfb*) and imaged at 11.5 hpf. Endoderm is outlined (dashed line), and endoderm width is indicated (solid line). Dorsal views, anterior towards the top. Scale bar, 150 μm. (C) Quantification of relative endoderm width (endoderm/embryo width) at 11.5 hpf from wild-type, heterozygous, and homozygous mutant *hgfa^fh528^* embryos injected with MO-*hgfb*. Points represent individual embryos color coded by batch. Open circles indicate batch means. Error bars indicate standard deviation. *p* values determined by one-way ANOVA. n.s., not significant. +/+ (n = 40), +/− (n = 59), −/− (n = 22), from four independent batches each. (D,D′) Endoderm labeled with *Tg(sox17:GFP)* from embryos injected with control (D) or *hgfa* plus *hgfb* (D′) morpholinos (MO-*hgfa/b*) and imaged at 11.5 hpf. Endoderm is outlined (dashed line), and endoderm width is indicated (solid line). Dorsal views, anterior towards the top. Scale bar, 150 μm. (E) Quantification of relative endoderm width (endoderm/embryo width) at 11.5 hpf from embryos injected with control or *hgfa/b* MO. Points represent individual embryos color coded by batch. Open circles indicate batch means. Error bars indicate standard deviation. *p* values determined by one-way ANOVA. n.s., not significant. MO-Ctrl (n = 85) and MO-*hgfa/b* (n = 84), from three independent batches each.

Although *hgfb* is not upregulated in *hgfa^fh528^* mutants, it remains possible that basal *hgfb* expression could compensate for the loss of *hgfa*. To test this, we used previously validated morpholino antisense oligonucleotides targeting *hgfa* and *hgfb* (MO-*hgfa* and MO-*hgfb*) (Anderson et al., 2013; Elsen et al., 2009). We first injected MO-*hgfb* into *hgfa^fh528^*embryos at the one-cell stage to eliminate both ligands. Embryos were imaged as before, and endoderm convergence was assessed by measuring relative endodermal width (Fig. 9B–B′′). Mean relative endodermal width was not significantly different between MO-*hgfb*-injected wild-type (0.473 ± 0.075), heterozygous (0.489 ± 0.066, *p* = 0.4285), and homozygous mutant (0.468 ± 0.068, *p* = 0.9398) *hgfa^fh528^* siblings (one-way ANOVA) (Fig. 9C). This result suggests that loss of both *hgfa* and *hgfb* does not impair endoderm convergence.

To address potential contributions from maternally deposited transcripts, we next performed simultaneous knockdown of *hgfa* and *hgfb* by co-injecting MO-*hgfa* and MO-*hgfb*. Morpholino efficacy was confirmed by impaired pancreas elongation at 3 dpf, consistent with prior reports (Anderson et al., 2013). Embryos were injected at the one-cell stage and imaged at 11 hpf, as described above. Mean relative endodermal width was not significantly different between control (0.518 ± 0.055) and MO-*hgfa/b*-injected embryos (0.530 ± 0.071, *p* = 0.2368, Student’s t-test) (Fig. 9 D–E).

Taken together, these results indicate that Hgf signaling is dispensable for endoderm convergence during gastrulation.

## DISCUSSION

Convergence of endoderm cells toward the midline requires a transition from dispersal to persistent, directional migration. While this behavior is often attributed to external signals, how endoderm cells themselves make this transition remains unclear.

Here, we report that the receptor tyrosine kinase Met promotes directional migration of endoderm cells during dorsal convergence. Although *met* is a marker of endoderm fate (Wan et al., 2026), its function in this cell type has not been examined in detail. We find that loss of Met delays endoderm convergence by reducing cell displacement and persistence, without affecting velocity, indicating that Met acts to enable efficient directional migration rather than controlling motility *per se*. Although Met is typically activated by Hgf, genetic evidence indicates that Hgf signaling is dispensable. Together, our findings identify Met as a regulator of migratory persistence during endoderm convergence and establish a ligand-independent role for Met signaling.

### A temporally distinct regulatory module in endoderm migration

Two features distinguish early *met* expression in the endoderm: its specificity and its timing. Unlike many genes involved in endoderm specification that are expressed at the onset of gastrulation, *met* expression becomes prominent during mid-gastrulation (Latimer and Jessen, 2008; Sur et al., 2023; Wagner et al., 2018). This timing sets it apart from early-acting transcription factors such as *sox32*, *sox17*, *gata5*, and *foxa2*, which are required for endoderm specification but are not endoderm-specific; they are also expressed in dorsal forerunner cells or mesodermal cell types, at least initially. In contrast, *met* is among the first genes to be expressed specifically in the endoderm in time to promote persistent, directional migration during convergence.

Two well-characterized regulators of endoderm migration, the chemokine receptor *cxcr4* and the Rac-GEF *prex1*, share this pattern: they are restricted to the endoderm, expressed during gastrulation, and control specific aspects of migratory behavior rather than the establishment of endoderm identity. Together, these observations suggest that *met* is not involved in fate specification but instead may be part of a distinct regulatory module that acts later to define endodermal cell behavior.

### Regulation of directional persistence in endoderm

Several signaling pathways regulate endoderm migration during gastrulation, acting at distinct stages and through different mechanisms (Nowotschin et al., 2019; Pinheiro and Heisenberg, 2020; Stainier, 2002; Zorn and Wells, 2009). Early in gastrulation, Nodal signaling, beyond its established role in specifying endoderm fate, promotes cell movement in part through activation of the Rac GEF *prex1* (Pézeron et al., 2008; Woo et al., 2012). In contrast, Bmp signaling contributes to dorsal convergence across the embryo, including in the endoderm. However, recent work indicates that Bmp elicits a comparatively weaker signaling response in endoderm than in mesoderm, and that perturbation of Bmp signaling specifically in endoderm has only modest effects on migration (Chang et al., 2025). These observations suggest that Bmp primarily influences endoderm behavior indirectly through its effects on mesoderm.

The Cxcl12–Cxcr4 systems coordinates endoderm and mesoderm movements. However, endodermal migration is not fully dependent on these external cues, as disruption of chemokine signaling delays convergence but does not abolish motility per se (Chang et al., 2025; Mizoguchi et al., 2008). This indicates that distinct features of endodermal cell behavior are differentially controlled and supports a model in which endoderm cells retain a degree of autonomy in their migratory behavior (Chang et al., 2025; Norris et al., 2017; Tavano et al., 2025). Consistent with this idea, we identify Met as a regulator that acts within the endoderm to promote efficient migration. Ultimately, loss of Met produces defects that closely resemble those observed upon disruption of Cxcr4 signaling—reduced displacement and persistence without major effects on instantaneous velocity (Chang et al., 2025; Mizoguchi et al., 2008). This points to shared regulation of migratory behavior by Met and Cxcl12–Cxcr4 signaling, which may act in parallel or converge on common downstream effectors (Cheng et al., 2018; Holland et al., 2013).

One potential point of convergence is Prex1. In neutrophils, Prex1 functions as a coincidence detector that requires dual inputs for activation, including GPCR signaling and phosphoinositide signaling (Welch et al., 2002). This dual-input requirement enables cells to integrate distinct extracellular cues into a unified migratory response. By analogy, Prex1 could integrate directional cues from chemokine signaling, mediated by the GPCR Cxcr4, with permissive or amplifying signals downstream of Met, which include activation of the PI3K–PIP3 pathway (Cheng et al., 2018; Organ and Tsao, 2011; Qiu et al., 2020; Trusolino et al., 2010). Notably, *prex1* is expressed earlier than both *met* and *cxcr4*, raising the possibility that it promotes general motility during dispersal, and directional migration later in gastrulation once Met and chemokine signaling become more prominent.

However, loss of either chemokine signaling (Chang et al., 2025; Mizoguchi et al., 2008) or Met (this study) attenuates but does not completely abolish migration directionality oriented toward the midline, indicating that directional migration is not solely dependent on these pathways and is likely supported by additional, partially redundant mechanisms. Met therefore promotes directional persistence, acting alongside additional pathways that support directional migration and convergence.

### Hgf-independence of Met signaling

In other developmental contexts, Met acts as a guidance cue receptor for Hgf, thus promoting directional migration (Elsen et al., 2009; Isabella et al., 2020; Nord et al., 2019). However, the expression patterns of the two *hgf* genes in zebrafish are not consistent with a role in regulating endoderm migration. *hgfb* is not expressed during gastrulation, while *hgfa* is activated towards the end of gastrulation, reaching appreciable levels after endoderm cells have become directional. Furthermore, Hgf is thought to diffuse poorly *in vivo*, thus limiting its range (Birchmeier et al., 2003). The spatially restricted expression of *hgfa* thus makes it unlikely to be a major driver of Met activation. Not surprisingly, our genetic data confirms that *hgfa* is dispensable.

This raises the question of how Met is activated in zebrafish endoderm in the absence of Hgf. Evidence for alternative mechanisms has been gathered in cancer models, where Met may be activated by cross-phosphorylation downstream of epidermal growth factor (Egf) signaling, or by extracellular matrix (ECM) engagement of integrins, causing Met activation through phosphorylation by integrin-linked kinase or its downstream targets (Jahangiri et al., 2017; Mitra et al., 2011; Stanislovas and Kermorgant, 2022), raising the possibility that similar noncanonical activation occurs during zebrafish gastrulation.

The zebrafish genome contains one Egf receptor, which is not expressed at significant levels in gastrula-stage endoderm (Sur et al., 2023); thus, Egf signaling is unlikely to strongly affect endoderm migration. In contrast, mesoderm and endoderm secrete integrin-dependent ECM proteins fibronectin and laminin during gastrulation (Latimer and Jessen, 2008). Concurrently, integrin receptors, such as integrin α5, become expressed in endoderm (Sur et al., 2023; Wagner et al., 2018), raising the possibility of coupling with Met signaling. Interestingly, loss of integrin α5 or fibronectin results in cardiac phenotypes (Gao et al., 2022); it is well-established that migrating cardiomyocytes use the endoderm as a substrate, and that defects in endoderm formation cause phenotypes, including pericardial edema (Yelon, 2001), similar to those we observed in Met-deficient larvae. Thus, integrin–ECM interaction may provide a mechanism for promoting Met-dependent directional migration.

Notably, in gastrointestinal cancers, Met signaling is primarily driven by elevated receptor expression rather than activating mutations (Bradley et al., 2017). Although Hgf may be present in the local environment and could contribute to receptor activation in some contexts, coordinated upregulation of ligand is unlikely to provide a robust or general mechanism for signaling. Consistent with this, elevated MET expression is strongly associated with poor patient prognosis in gastrointestinal cancers (Miekus, 2017). Our data therefore suggest that the ligand-independent activity we observe in vivo may reflect a broader mode of Met regulation relevant to both development and disease.

## MATERIAL AND METHODS

### Zebrafish

Adult zebrafish were maintained under standard laboratory conditions. All embryos used in this study were of an early stage prior to sex identification. Embryos were raised in egg water at 28.5°C. Zebrafish in an outbred background (Ab/Tl/Ekw) were used as wild type. *Tg(sox17:GFP)^s870^* (Chung and Stainier, 2008), *met^fh533^*, and *hgfa^fh528^* have been previously described (Isabella et al., 2020). *met^ucm123^* and *met^ucm124^* were generated for this study (see below). This study was performed with the approval of the Institutional Animal Care and Use Committee (IACUC) of the University of California Merced (Protocol #2023–1144).

### Generation of *met^ucm123^ and met^ucm124^* mutant zebrafish

We generated transcription start site (tss) deletions of the *met* locus using CRISPR/Cas9 mutagenesis as previously described (Zhao et al., 2021). Two guide RNAs (gRNAs) were chosen based on “high predicted cleavage” as determined by the CRISPR tracks of the UCSC Genome Browser (Casper et al., 2026). One gRNA (5′-TAAATGATCCGTCGTCAATG-3′) targeted upstream of the tss as determined by the RNA sequencing tracks of the UCSC Genome Browser. The other gRNA (5′-TCATTCAGAAGGGTTACACA-3′) targeted the intron between exons 2 and 3. Guide RNAs were synthesized as Alt-R™ CRISPR-Cas9 sgRNAs by Integrated DNA Technologies (IDT). *Tg(sox17:GFP)* embryos were injected at the 1-cell stage with 100 pg gRNA and Alt-R™Cas9 mRNA as per manufacturers recommendations.

For founder screening, injected F_0_ fish were raised to adulthood then outcrossed to wild-type zebrafish. From each clutch, 35 embryos were pooled at 24 hpf and genomic DNA was isolated as previously described (Meeker et al., 2007). The *met* locus was amplified by primers flanking the gRNA target sites (Fwd, 5’-TGGTAAACTATCATTTCTGTTGTTGGTCTG-3’; Rev: 5’-TAGGGTATAAAGACAAGCAATTGGGTTGAC-3’) using GoTaq G2 Green Master Mix with the following program: 95 °C for 5 min, followed by 35 cycles of 95 °C for 30 s, 56 °C for 30 s, and 72 °C for 45 s. PCR fragments were cloned into pGEM-T Easy (Promega), and the inserts were sequenced. *met^ucm123^* harbors a 5918 bp deletion and *met^ucm124^* a 5914 bp deletion. Only clutches containing the expected deletion were kept for propagation. At adulthood, individual F_1_ zebrafish were genotyped by fin clipping using the same primer sets as described above. Only animals containing identical deletion mutations were kept for line propagation.

Genotyping was performed using two sets of primers: 1) the deletion-flanking primers described above , which produce a 363 bp product from *met^ucm123^*, a 367 bp product from *met^ucm124^*, and no product from wild type, and 2) primers flanking exon 2 (F: 5’-TTTGAGTGCATTTACTCTGAGCGAAGGAGA-3’; R: 5’-TAGGGTATAAAGACAAGCAATTGGGTTGAC-3’) which produce a 430 bp product from wild type and no product from *met^ucm123^* or *met^ucm124^*.

### Met inhibitor treatments

For pharmacological inhibition of Met activity, embryos were treated with 30 mM INCB28060(Liu et al., 2011), 50 mM SGX-523 (Buchanan et al., 2009), or 1% dimethyl sulfoxide (DMSO; Sigma-Aldrich) as a vehicle control starting at 3.5 hp. Chorions were punctured prior to treatment.

### Morpholino-mediated knockdown

We performed knockdown of *hgfa* and *hgfb* using previously reported morpholino antisense oligonucleotides (Anderson et al., 2013; Elsen et al., 2009): MO2-*hgfa* (ZFIN_ID: ZDB-MRPHLNO-100420-4); MO1-*hgfb* (ZFIN_ID: ZDB-MRPHLNO-100420-3). As a control, we used a previously described standard control morpholino (5’-CCTCTTACCTCAGTTACAATTTATA -3’; Kim et al., 2007; Paraiso et al., 2019). Morpholinos were synthesized by Gene Tools, LLC; 1 ng of morpholino was injected at the one-cell stage. To quantify *met*, *hgfa*, and *hgfb* transcript levels, total RNA was extracted from 35 embryos using the RNeasy Micro Kit (Qiagen). For single-embryo qPCR, DNA and RNA were extracted using the AllPrep DNA/RNA Micro Kit (Qiagen). To ensure proper lysis, embryos were homogenized by aspiration through a 31G ultra-fine needle. cDNA was synthesized from 1 µg of total RNA using the ProtoScript II First Strand cDNA Synthesis Kit (New England Biolabs). Each qPCR reaction contained 2X PerfeCTa® SYBR green fast mix (Quantabio), 5-fold diluted cDNA and 325 nM each primer. Reactions were performed in a QuantStudio 3 real-time PCR machine (Applied Biosystems) using the following program: 95 °C for 10 min, followed by 40 cycles of 95 °C for 30 s, 60 °C for 30 s, and 72 °C for 1 min. Melting curve analysis was performed to confirm specificity. Data represent averages from three biological replicates, each with three technical replicates. Relative expression levels were calculated using the 2^−ΔΔCT^ method. The housekeeping gene *eef1a1l1* was used as an internal reference (Ligunas and Materna, 2026). For absolute quantification and comparison across developmental stages, an external reference was used, as previously described (Ligunas and Materna, 2026). Primer sequences were as follows (5′ to 3′; e, exon):

*hgfa* forward: ATGGATGGGGAGACACTAAAGGACA

*hgfa* reverse: TTTCTCACAAACGCCCTGATCTCTC

*hgfb* forward: AAACCAAAGGCACTGGACATGAGG

*hgfb* reverse: CGTAGTCTTTCTCACAAACTCCTTC

*met*-e9 forward: CCCAAACATTCTTTCATCAGTGGAGG

*met*-e11 reverse: TCATCATGACTGCAGGTCACTTGGAA

*met*-e18 forward: AGAAACTGCATGCTGGATGAGAGC

*met*-e20 reverse: AAAACACCAAACGACCACACATCCG

eef1a1l1 forward: CACGGTGACAACATGCTGGAG

eef1a1l1 reverse: CAAGAAGAGTAGTACCGCTAGCAT

### Hybridization chain reaction RNA-fluorescent *in situ* hybridization (HCR RNA-FISH)

Whole mount in situ hybridization analysis of *met* and *hgfa* expression was performed by HCR RNA-FISH as previously described (Ibarra-García-Padilla et al., 2021) using *Tg(sox17:GFP)* embryos; GFP fluorescence survived fixation and processing, allowing for identification of endodermal cells without the need for additional labeling. Briefly, embryos at the indicated stages were fixed in 4% paraformaldehyde overnight at 4°C, rinsed with phosphate-buffered saline (PBS), and dehydrated in 100% methanol overnight at -20°C. Prior to hybridization, embryos were sequentially rehydrated with methanol in PBST. HCR RNA-FISH amplifiers, buffers and probes against zebrafish *met* (XM_005162995.4) and *hgfa* (XM_005164742.3) were obtained from Molecular Instruments.

### Microscopy, image analysis, and image processing

Spinning disk confocal imaging was performed as previously described (LaBelle et al., 2025). For live imaging, embryos were manually dechorionated and embedded in 1% low-melting agarose in 35 mm glass-bottom dishes (MatTek Corporation) and submerged in 0.3X Ringer’s solution. For Met inhibitor treatment experiments, embryos were submerged in 0.3X Ringer’s solution containing 1% DMSO alone or with Met inhibitor. For HCR RNA-FISH imaging, embryos were incubated in 5X saline-sodium citrate buffer plus 0.1% Tween-20 (SSCT) supplemented with 30% glycerol and flat mounted on glass slides as previously described (Cheng et al., 2014).

Images were acquired with a 10x/0.4 NA or 30x/1.05 NA objective lens on a microscope (IX83; Olympus) equipped with a spinning disk confocal unit (CSU-W1; Andor), a scientific complementary metal–oxide–semiconductor (sCMOS) camera (Prime 95b; Teledyne Photometrics), and MicroManager software 78. The microscope stage was enclosed in a temperature-controlled case, and live samples were kept at 28.5°C. Z-stacks were acquired at 2 µm intervals. For time lapse imaging, Z-stacks were acquired every 5 minutes. For HCR RNA-FISH imaging, multiple tiled Z-stacks were acquired to capture the entire length of each embryo.

Image analysis and processing was performed using Fiji software (Schindelin et al., 2012). All measurements and subsequent processing were performed from maximum projections of confocal Z-stacks. Average migration displacement, instantaneous velocity, and persistence were measured using the TrackMate plugin for ImageJ (Tinevez et al., 2017); at least 3 embryos were analyzed per condition. For tiled images, maximum Z-projections were stitched together using the pairwise stitching function in Fiji. For all figures and videos, images were processed in Fiji as follows: background subtracted, brightness levels adjusted, converted to 8-bit depth, and cropped.

### Statistical analysis

Plots and statistical analyses were performed using Prism (GraphPad). For inhibitor treatments, *p* values were determined using Student’s t-test. Cell migration parameters (velocity, displacement, persistence) were quantified for individual cells using TrackMate. For statistical analysis, values were averaged per embryo, and these embryo-level means were treated as independent biological replicates; *p* values were determined using one-way ANOVA. Unless otherwise noted, data are reported as mean ± standard deviation (SD).

## ACKNOWLEDGEMENTS

We would like to thank Cecilia Moens for sharing *met^fh533^* and *hgfa^fh528^* zebrafish lines. We are grateful for excellent animal care by the Department of Animal Research Services. We are thankful for many insights and suggestions from members of the Woo and Materna labs, especially Gloria Ligunas and Jesselynn LaBelle. This work was supported by grants from National Institutes of Health (R21HD107313) and the National Science Foundation (IOS-2238304) to SW and startup funds to SCM.

## FIGURE LEGENDS

**Video 1. Time-lapse video of Met inhibitor-treated embryo during endoderm convergence.** Migrating endodermal cells are GFP-labeled in *Tg(sox17:GFP)*. Embryos were treated with control vehicle (DMSO) or Met inhibitor (INCB28060) prior to gastrulation (3.5 hpf) and imaged starting at 8.5 hpf over a ∼1.5 hour window that encompasses dorsal convergence of endoderm. Embryos viewed laterally with dorsal toward the right and time displayed as min (top left corner), related to Fig. 3.

**Video 2. Time-lapse video of *met^ucm124^* embryos during endoderm convergence.** Migrating endodermal cells are GFP-labeled in *Tg(sox17:GFP)*. Wild-type (+/+), heterozygous (+/-), or homozygous (-/-) *met^ucm124^* embryos were obtained by intercross of *met^ucm124/+^*parents, followed by live-imaging starting at 8.5 hpf over a ∼1.5 hour window that encompasses dorsal convergence of endoderm. Embryos viewed laterally with dorsal toward the right and time displayed as min (top left corner), related to Fig. 6.

**Video 3. Time-lapse video of *met^fh533^* embryos during endoderm convergence.** Migrating endodermal cells are GFP-labeled in *Tg(sox17:GFP)*. Wild-type (+/+), heterozygous (+/-), or homozygous (-/-) *met^fh533^* embryos were obtained by intercross of *met^fh533/+^*parents, followed by live-imaging starting at 8.5 hpf over a ∼1.5 hour window that encompasses dorsal convergence of endoderm. Embryos viewed laterally with dorsal toward the right and time displayed as min (top left corner), related to Fig. 6.

**Video 4. Time-lapse video of *hgfa^fh528^* embryos during endoderm convergence.** Migrating endodermal cells are GFP-labeled in *Tg(sox17:GFP)*. Wild-type (+/+), heterozygous (+/-), or homozygous (-/-) *met^fh528^* embryos were obtained by intercross of *met^fh528/+^*parents, followed by live-imaging starting at 8.5 hpf over a ∼1.5 hour window that encompasses dorsal convergence of endoderm. Embryos viewed laterally with dorsal toward the right and time displayed as min (top left corner), related to Fig. 8.

## Declaration of generative AI and AI-assisted technologies in the manuscript preparation process

During the preparation of this work, the authors used ChatGPT in order to assist with language editing, phrasing, and clarity of the manuscript text. After using this tool, the authors reviewed and edited the content as needed and take full responsibility for the content of the published article.

---

**Supplemental Figure 1.**
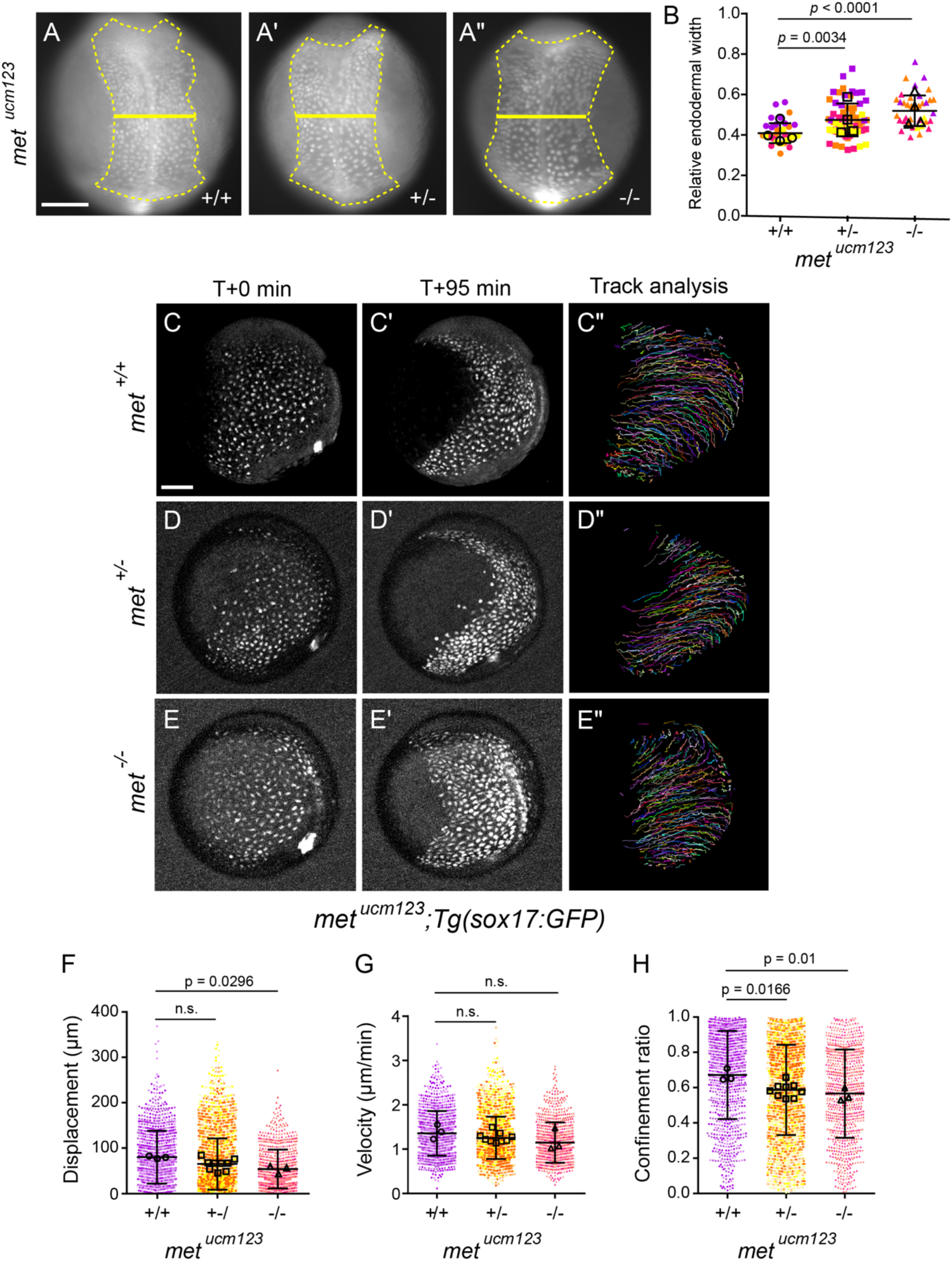
m*e*t^ucm123^ exhibits delayed endoderm convergence and reduced migration persistence. (A–A′′) Representative images of endoderm labeled with *Tg(sox17:GFP)* in wild-type (A), heterozygous (A′), and homozygous mutant (A′′) *met^ucm123^* embryos at 11.5 hpf. Endoderm is outlined (dashed line), and endoderm width is indicated (solid line). Dorsal views, anterior towards the top. Scale bar, 150 μm (B) Quantification of relative endoderm width (endoderm/embryo width) at 11.5 hpf from wild-type, heterozygous, and homozygous mutant *met^ucm123^* embryos at 11.5 hpf. Points represent individual embryos color coded by batch. Open circles indicate batch means. Error bars indicate standard deviation. *p* values determined by one-way ANOVA. +/+ (n = 32), +/− (n = 68), −/− (n = 37), from four independent batches each. (C–E′′) Time-lapse confocal imaging of endoderm labeled with *Tg(sox17:GFP)* in wild-type (C–C′′), heterozygous (D–D′′), and homozygous mutant (E–E′′) *met^ucm123^* embryos starting at 8.5 hpf. Embryos are shown at the start (C, D, E) and end (C′, D′, E′) of each time-lapse along with migration tracks generated by cells throughout the time-lapse (C′′, D′′, E′′). Lateral views, dorsal to the right. Scale bar, 100 μm. (F–H) Quantification of migration displacement (F), velocity (G), and persistence (H) from wild-type, heterozygous, and homozygous mutant *met^ucm123^*embryos. Points represent individual cell tracks, color colored by embryo. Open circles depict embryo means. Error bars indicate standard deviation. *p* values determined by one-way ANOVA. n.s., not significant. +/+ (987 cells from 3 embryos), +/− (2835 cells from 9 embryos), −/− (832 cells from 3 embryos).

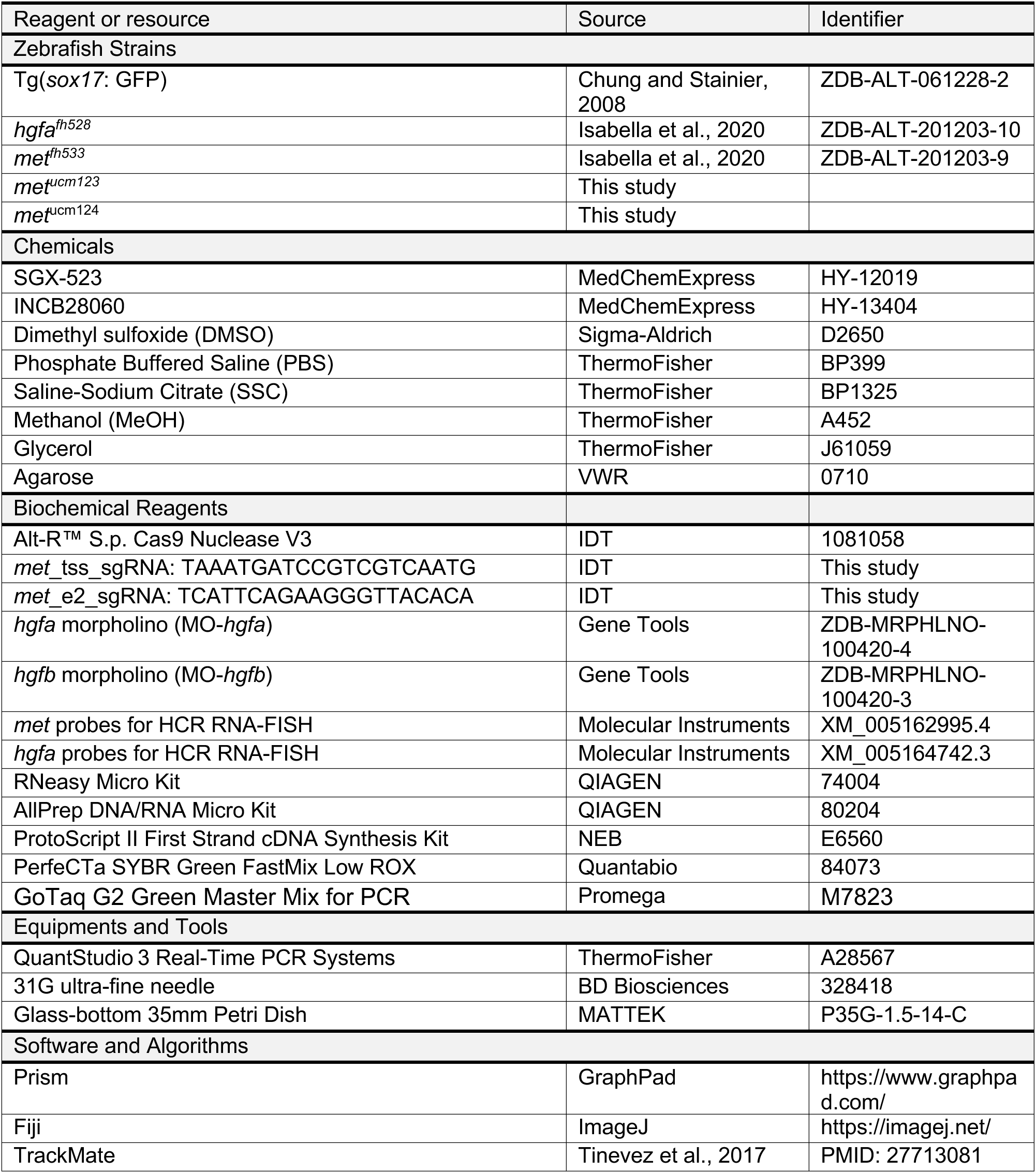

